# Nanograms of SARS-CoV-2 Spike Protein Delivered by Exosomes Induce Potent Neutralization of Both Delta and Omicron Variants

**DOI:** 10.1101/2023.03.20.533560

**Authors:** Mafalda Cacciottolo, Yujia Li, Justin B Nice, Michael J. LeClaire, Ryan Twaddle, Ciana L. Mora, Stephanie Y. Adachi, Meredith Young, Jenna Angeles, Kristi Elliott, Minghao Sun

## Abstract

Exosomes are emerging as potent and safe delivery carriers for use in vaccinology and therapeutics. A better vaccine for severe acute respiratory syndrome coronavirus 2 (SARS-CoV-2) is needed to provide improved, broader, longer lasting neutralization of SARS-CoV-2, a more robust T cell response, enable widespread global usage, and further enhance the safety profile of vaccines given the likelihood of repeated booster vaccinations. Here, we use Capricor’s StealthX™ platform to engineer exosomes to express native SARS-CoV-2 spike Delta variant (STX-S) protein on the surface for the delivery of a protein-based vaccine for immunization against SARS-CoV-2 infection. The STX-S vaccine induced a strong immunization with the production of a potent humoral immune response as demonstrated by high levels of neutralizing antibody not only against the delta SARS-CoV-2 virus but also two Omicron variants (BA.1 and BA.5), providing broader protection than current mRNA vaccines. Additionally, both CD4^+^ and CD8^+^ T cell responses were increased significantly after treatment. Quantification of spike protein by ELISA showed that only nanograms of protein were needed to induce a potent immune response. This is a significantly lower dose than traditional recombinant protein vaccines with no adjuvant required, which makes the StealthX™ exosome platform ideal for the development of multivalent vaccines with a better safety profile. Importantly, our exosome platform allows novel proteins, or variants in the case of SARS-CoV-2, to be engineered onto the surface of exosomes in a matter of weeks, comparable with mRNA vaccine technology, but without the cold storage requirements. The ability to utilize exosomes for cellular delivery of proteins, as demonstrated by STX-S, has enormous potential to revolutionize vaccinology by rapidly facilitating antigen presentation at an extremely low dose resulting in a potent, broad antibody response.

## IINTRODUCTION

The emergence of severe acute respiratory syndrome coronavirus 2 (SARS-CoV-2) has created an urgent need for vaccine development strategies to produce safe, effective, readily available and accessible vaccines that can be efficiently produced to combat the emergence of evolving SARS-CoV-2 variants [1] and any future outbreaks. Many vaccines against SARS-CoV-2 have been developed; all candidates try to achieve immunity to the virus, mainly directing the immune response to surface spike protein, which binds to the host cell receptor angiotensin-converting enzyme 2 (ACE2), mediating viral cell entry and infection[2]. The spike (S) protein is a class I fusion glycoprotein, the major surface protein on the SARS-CoV-2 virus, which mediates binding to the ACE receptor on the cell surface promoting its propagation and infection. It is the primary target for neutralizing antibodies. Spike is also the primary site of mutations identified in SARS-CoV-2 virus, potentially reducing the efficacy of the immune-response induced by certain vaccines. Since the beginning of the COVID-19 pandemic, five variants of concern (VOC) and eight variants of interest (VOI) have been reported by the World Health Organization, posing challenges to current vaccines, and creating the need for more effective vaccines.

Current leading SARS-CoV-2 vaccines have employed mRNA-lipid nanoparticle and recombinant proteins [3]. Recombinant protein vaccines are a safe and reliable approach for immunization, but generally suffer from weak immunogenicity, thus requiring formulation with an appropriate adjuvant to achieve a strong immune response [4, 5]. Additionally, recombinant protein vaccines suffer from long production times, limiting rapid response to new variants. While all current mRNA vaccine candidates achieve an initial immune response to the virus with consequent reduction of hospitalization rates, they all lack long-term, multi-variant protection. It has been reported that antibody levels drastically decline over time and therefore the efficacy to protect against infection drastically declines as well [6]. Moreover, mRNA vaccines have lacked cross-reactivity against new VOC, requiring multiple booster injections to maintain protection against the virus. Thus, a better, globally deployable vaccine for SARS-CoV-2 is needed to provide improved, broader, longer lasting neutralization of SARS-CoV-2, a more robust T cell response, and further enhance the safety profile of vaccines [7].

Exosomes were first identified in 1983 by Stahl [8] and Johnstone [9] as extracellular membrane vesicles with unknown function. Subsequently, many studies have contributed to identifying exosomes’ role in physiology and disease through cell-to-cell communication. Exosomes are small vesicles of ∼30-200 nm diameter, enriched in specific subsets of proteins (CD81, CD63, CD9; Tetraspanin family), RNAs, and lipids [10]. Recently, utilization of exosomes as a delivery system has garnered great interest. Reduced toxicity together with exosomes’ natural role in cell communication and the endless opportunities for exosome engineering for cell-specific targeting and vaccinology make exosomes an excellent delivery system. Importantly, isolation of exosomes from different cell types, including 293F cells [11] is established and amenable to scale up purification processes. In addition, 293F exosomes do not carry class I and class II major histocompatibility (MHC) proteins and B7 co-stimulatory molecules which are involved in immune response stimulation; thus, their absence on exosomes allows for allogeneic use and ensures safety [12]. Moreover, exosomes can efficiently deliver antigens to immune cells due to the presence of several adhesion proteins on their surface [13] ; thus, exosomes can present foreign antigens efficiently to antigen presenting cells such as dendritic cells, which present the antigen to T cells for facilitating antibody production and cytotoxic activity.

Here, we developed a SARS-CoV-2 vaccine by utilizing our exosome platform (StealthX™, STX) to engineer exosomes to express the native SARS-CoV-2 Delta spike protein (STX-S) on their surface. STX-S exosomes were used to deliver a protein-based vaccine for prevention and immunization against SARS-CoV-2 infection in mice. These data show that the STX vaccine elicits both CD4^+^ and CD8^+^ T cell responses and induces a potent humoral immune response as demonstrated by neutralizing antibody not only against the delta SARS-CoV-2 virus but also Omicron variants. Importantly, the robust immune response observed after STX vaccination was induced using only nanogram amounts of SARS-CoV-2 protein displayed on exosomes in absence of any adjuvant or lipid nanoparticle, which further strengthens the STX exosome safety profile as a vaccine candidate. Additionally, due to the significantly lower amount of protein used for vaccination, multivalent vaccines are easily generated using the STX platform.

These results support the clinical development of the STX-S vaccine for immunization against COVID-19.

## METHODS

### Cell lines

Human embryonic kidney 293 T cells (293T) were purchased from ATCC (CRL-3216). 293T cells were maintained in culture using Dulbecco’s Modified Eagle Medium (DMEM), high glucose, Glutamax™ containing 10% fetal bovine serum. 293T cells were incubated at 37℃ /5% CO_2_. FreeStyle™ 293F cells (Gibco, 51-0029) were purchased from ThermoFisher. 293F cells were used as a parental cell line to generate spike SARS-CoV-2 delta spike expressing stable cell lines: Stealth X-Spike cells (STX-S). 293F and STX-S cells were maintained in a Multitron incubator (Infors HT) at 37℃, 80% humidified atmosphere with 8% CO_2_ on an orbital shaker platform rotating at 110 rpm.

### Lentiviral particle and STX-S cell line generation

Lentiviral vectors for expression of SARS-CoV-2 spike (Delta variant B.1.617.2, NCBI accession # OX014251.1) were designed and synthesized from Genscript together with the two packaging plasmids (pMD2.G and psPAX2). Lentiviral particles for spike transduction were generated by transfecting 293T cells with pMD2.G (Genscript), psPAX2 (Genscript) and STX-S_pLenti at a ratio of 5:5:1 using Lipofectamine 3000 (ThermoFisher, L3000008) according to the manufacturer’s instruction. Spike lentiviral particles were collected at 72 hours (h) post transfection and used to transduce 293F parental cells to generate STX-S cells.

### Flow cytometry

Standard flow cytometry methods were applied to measure the spike SARS-CoV-2 protein expression on STX-S cell surface. In brief, 250K STX-S cells were aliquoted, pelleted and resuspended in 100µL eBioscience™ Flow Cytometry Staining Buffer (ThermoFisher, 00-4222-57). Cells were incubated at room temperature (RT) for 30 min protected from light in the presence of anti-spike antibody (Abcam, clone 1A9, ab273433) labeled with Alexa Fluor®-647 (Alexa Fluor® 647 Conjugation Kit (Fast)-Lightning-Link® (Abcam, ab269823) according to the manufacturer’s protocol. Following incubation, STX-S cells were washed with eBioscience™ Flow Cytometry Staining Buffer (ThermoFisher, cat No 00-4222-57), resuspended in PBS and analyzed on the CytoFlex S flow cytometer (Beckman Coulter). Data was analyzed by FlowJo (Becton, Dickinson and Company; 2021).

### Cell sorting

Cell sorting was performed at the Flow Cytometry Facility at the Scripps Research Institute (San Diego, CA). To enrich the spike positive population, STX-S cells were stained as described above for flow cytometry and went through cell sorting (Beckman Coulter MoFlo Astrios EQ) to generate pooled STX-S. The pooled STX-S were used in all the experiments in this paper unless specified otherwise.

### STX-S Exosome Production

STX-S cells were cultured in FreeStyle media (ThermoFisher, 12338018) in a Multitron incubator (Infors HT) at 37℃, 80% humidified atmosphere with 8% CO_2_ on an orbital shaker platform. Subsequently, cells and cell debris were removed by centrifugation, while microvesicles and other extracellular vesicles larger than ∼220 nm were removed by vacuum filtration. Next, exosomes were isolated using either Capricor’s lab scale or large-scale purification method. For lab scale: supernatant was subjected to concentrating filtration against a Centricon Plus-70 Centrifugal filter unit (Millipore, UFC710008), then subjected to size exclusion chromatography (SEC) using a qEV original SEC column (Izon, SP5). For large scale: supernatant was subjected to concentrating tangential flow filtration (TFF) on an AKTA Flux s instrument (Cytiva, United States) and then subjected to chromatography on an AKTA Avant 25 (Cytiva, United States).

### Nanoparticle Tracking Analysis

Exosome size distribution and concentration were determined using ZetaView Nanoparticle Tracking Analysis (Particle Metrix, Germany) according to manufacturer instructions. Exosome samples were diluted in 0.1 µm filtered 1X PBS (Gibco, 10010072) to fall within the instrument’s optimal operating range.

### Jess Automated Western Blot

Detection of SARS-CoV-2 spike proteins in cell lysate and exosomes used Protein Simple’s Jess capillary protein detection system. Samples were lysed in RIPA buffer (ThermoFisher Scientific, 8990) supplemented with protease/phosphatase inhibitor (ThermoFisher Scientific, A32961), quantified using the BCA assay (ThermoFisher Scientific, 23227) and run for detection. To detect spike, the separation module 12–230 kDa was used following manufacturers protocol. Briefly, 0.8 µg of sample and protein standard were run in each capillary, probed with anti-mouse Ms-RD-SARS-COV-2 (R&D Systems, MAB105401) followed by HRP secondary antibody provided in Jess kits (Protein Simple, 042-205).

### TEM imaging for characterization of STX-S exosome morphology

STX-S exosome samples were negatively stained onto copper grids with a carbon/Formvar film coating and imaged by TEM at the Electron Microscopy Core Facility at UC San Diego (San Diego, CA). Briefly, samples were treated with glow discharge, stained with 2% uranyl acetate and dried before imaging. Grids were imaged on a JEM-1400 Plus (JEOL Ltd, Japan) at 80kV and 48µA. Images were taken at from 12K-80K magnification at 4kx4k pixels of resolution.

### CD81 bead-assay

STX-S or 293F parental exosomes were mixed with anti-CD81 labeled magnetic beads overnight at 4°C (ThermoFisher, 10622D) and washed twice with PBS using the magnetic stand. Next, the bead-exosomes were incubated with either direct conjugated AlexaFluor 647 anti-spike (see above flow cytometry), FITC anti-CD81 antibody (BD Biosciences, 551108), or FITC Mouse IgG, κ isotype control (BD Biosciences, 555748) for 1 h at RT followed by PBS wash. 293F exosomes were used as a negative control for spike expression and the isotype antibody was used as a negative control for CD81 expression. Samples were analyzed on a CytoFlex S (Beckman Coulter) flow cytometer and data was analyzed by FlowJo.

### Animal Studies

To examine the efficacy of STX-S exosomes, age matched BALB/c mice (female, 8-10 wks old) were anesthetized using isoflurane and received bilateral intramuscular injection (50 µl per leg, total 100 µl) of 1) PBS or 2) STX-S exosomes. Four doses were used throughout the study: STX-S dose 1 = 0.32 ng, STX-S dose 2 = 3 ng, STX-S dose 3 = 10 ng, STX-S dose 4 = 33 ng. Booster injection was performed at day 21. Mice were monitored closely for changes in health and weight was recorded biweekly. Blood collection was performed at day 14 and day 35. Blood was collected from the submandibular vein and processed for plasma isolation after centrifugation. For comparison study, mice were injected with equal amounts of SARS-CoV-2 protein delivered as 1) soluble protein in conjugation with adjuvant (Alhydrogel, 100 µg/dose, vac-alu-250, InvivoGen), or 2) STX-S exosome. Blood was collected 2 weeks after injection and tested for IgG against SARS-CoV-2. Timeline of studies is outlined in Fig.1.

**Figure 1.**
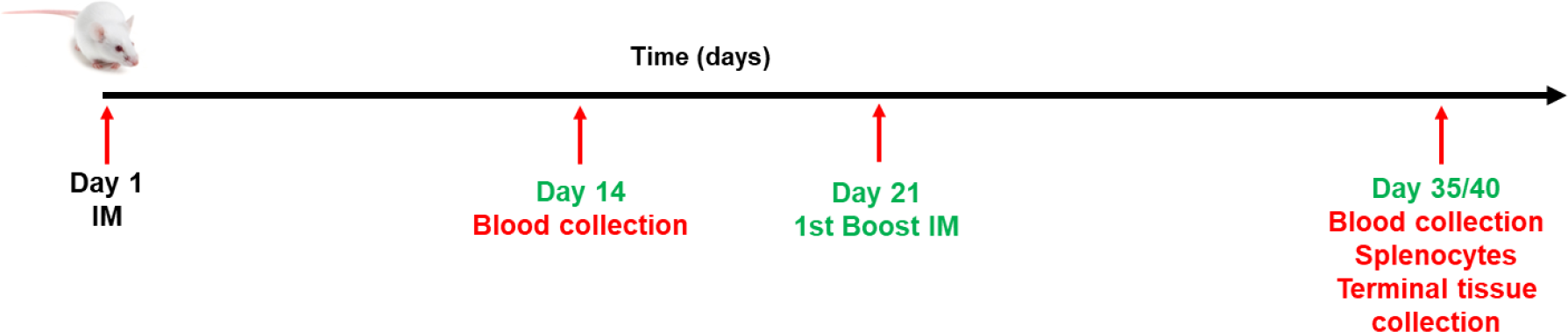
Schematic diagram of in vivo studies.

### IgG ELISA

Mouse IgG antibody against SARS-CoV-2 spike was measured by enzyme-linked immunosorbent assays (ELISA) using precoated ELISA plates (IEQ-CoV-S-RBD-IgG, RayBiotech) according to the manufacturer’s instructions, at RT. Briefly, mouse plasma samples were diluted in sample buffer (RayBiotech) and added to antigen-coated wells, in duplicates, and incubated at RT for 2 h on a shaker. Commercially available antibody against Spike (S1N-S58, Acro Biosystems) was used as positive control. Plates were washed, incubated for 1 h at RT with HRP-conjugated goat anti-mouse secondary antibodies (111-035-003, Jackson ImmunoResearch) diluted in assay buffer (RayBiotech). After washing, plates were developed using TMB substrate (RayBiotech). The reaction was stopped by adding STOP solution and the absorbance at 450 nm was recorded using a BioTeck Gen5 plate reader (Agilent). Endpoint titers were calculated as the dilution that emitted an optical density exceeding 4X the PBS control group.

### Neutralizing antibodies against DELTA SARS-CoV-2

Samples were analyzed by Retrovirox, Inc. (San Diego, CA). Briefly, Vero E6 cells were used to evaluate the neutralization activity of the test-items against a replication competent SARS-CoV-2 delta variant (B.1.617.2). Samples were pre-incubated with the virus for 1 h at 37°C before addition to cells. Following pre-incubation of plasma/virus samples, cells were challenged with the mixture. Samples were present in the cell culture for the duration of the infection (96 h), at which time a *Neutral Red* uptake assay was performed to determine the extent of the virus-induced cytopathic effect (CPE). Prevention of the virus-induced CPE was used as a surrogate marker to determine the neutralization activity of the test-items against SARS-CoV-2. Test-items were evaluated in duplicates using two-fold serial dilutions starting at a 1:40 dilution (eight total dilutions). Control wells included “CoV02-Delta” and GS-441524, tested in singlet data points on each plate. “CoV02-Delta” is convalescent plasma from an individual who previously received two doses of the Moderna COVID-19 vaccine before infection with the Delta variant. GS-441524 is an antiviral from Gilead Sciences. NT50 values of the test-items were determined using *GraphPad Prism* software.

### Neutralizing antibodies against Omicron (BA.1 and BA.5.2.1) SARS-CoV-2

Samples were analyzed by Retrovirox, Inc. (San Diego, CA). Neutralization assays were performed using anti-NP immunostaining (Omicron BA.1 and BA.5.2.1). Briefly, samples were pre-incubated with the virus for 1 h at 37°C before addition to Vero E6 cells. Following incubation, media was removed, and then cells were challenged with the SARS-CoV-2 / test-item pre-incubated mix. The amount of viral inoculum was previously titrated to result in a linear response inhibited by antivirals with known activity against SARS-CoV-2. Cell culture media with the virus inoculum was not removed after virus adsorption, and test-items and virus were maintained in the media for the duration of the assay (48 h). Subsequently, the extent of infection is monitored by incubating cells with a monoclonal test-item against the SARS-CoV-2 nucleocapsid (NP). The amounts of the viral antigen in infected cells are estimated after incubation with horseradish peroxidase conjugated polyclonal test-items against human IgG (HRP-goat anti-mouse IgG). The reaction is monitored using a colorimetric readout (absorbance at 492nm). Test-items were evaluated in duplicates using two-fold serial dilutions starting at a 1:40 dilution. Control wells included GS-441524 (Gilead Sciences), tested in singlet data points on each plate.

### Splenocyte isolation

Spleens were processed for single cell isolation by mechanical disruption of the spleen pouch using a syringe stopper and passed through a nylon cell strainer to remove tissue debris. Erythrocytes were lysed using ammonium chloride potassium (ACK) buffer (A1049201, ThermoFisher) and splenocytes were collected by centrifugation. Cellular pellet was resuspended in completed RPMI 1640 media (FG1215, Millipore Sigma Aldrich).

### ELISPOT

Briefly, splenocytes were isolated by mechanical disruption of spleen pouch and seeded at a concentration of 5E5 cells/well in a precoated 96-well plate and incubated for 24 h in the presence or absence of 10 µg/ml of SARS-CoV-2 Spike (S1N-C52H4, AcroBiosystems). Commercially available ELISPOT plates for evaluation of IL-4 (MuIL4, Immunospot, Cellular Technology Limited), and IFNγ (MuIFNγ, Immunospot, Cellular Technology Limited) were used. Assay was performed according to manufacturer’s guidelines. Plates were analyzed using the ELISPOT reader S6ENTRY (Immunospot, Cellular Technology Limited).

### ELISA for protein quantification

Spike protein level on exosomes was measured by ELISA using precoated ELISA plates (ELV-COVID19S1, RayBiotech) according to the manufacturer’s instructions, at RT. Briefly, samples and standards were loaded onto the precoated plate, and incubated on a shaker. Plates were washed and incubated with biotin-conjugated detection antibody for an hour at RT, followed by 45 min incubation in streptavidin solution. After washes, plates were developed for 30 min using TMB substrate. The reaction was stopped by adding STOP solution and the absorbance at 450 nm using a BioTeck Gen5 plate reader (Agilent).

### Complete Blood count

Fresh plasma at the time of terminal collection was analyzed according to the manufacturer’s instructions using the Hemavet 950FS (Drew Scientific).

### Pathology

Tissues (brain, salivary gland, heart, lung, liver, spleen, kidney, gastro-intestinal tract (GI) and skeletal muscle (site of injection)) were collected and fixed in fixed in 10% Neutralized Formalin. Tissues were sent to Reveal Biosciences (San Diego, CA) for further processing and analysis. Sections were stained with hematoxylin and eosin and analyzed for alterations. Pathology evaluation was performed by Dr Mary E.P. Goad (DVM, PhD, DACVP, DACLAM, DABT).

### Statistical analysis

Data were analyzed using Excel and GraphPad Prism 9.1 and shown as mean±sem. 1-way ANOVA with post-hoc correction for multiple comparisons or 2-tailed t-test were applied as needed.

## RESULTS

### SARS-CoV-2 spike expression on the surface of STX-S cells and exosomes

To facilitate expression of naïve SARS-CoV-2 delta spike (STX-S) on the exosomes, STX-S cells were generated to express SARS-CoV-2 spike on their surface by lentiviral transduction of 293F cells. Expression of SARS-CoV-2 spike on the cell surface was evaluated by flow cytometry. As shown in Figure 2A, >97% of STX-S cells expressed spike on their surface. Non-engineered, parental 293F cells do not express spike, as expected (data not shown).

**Figure 2.**
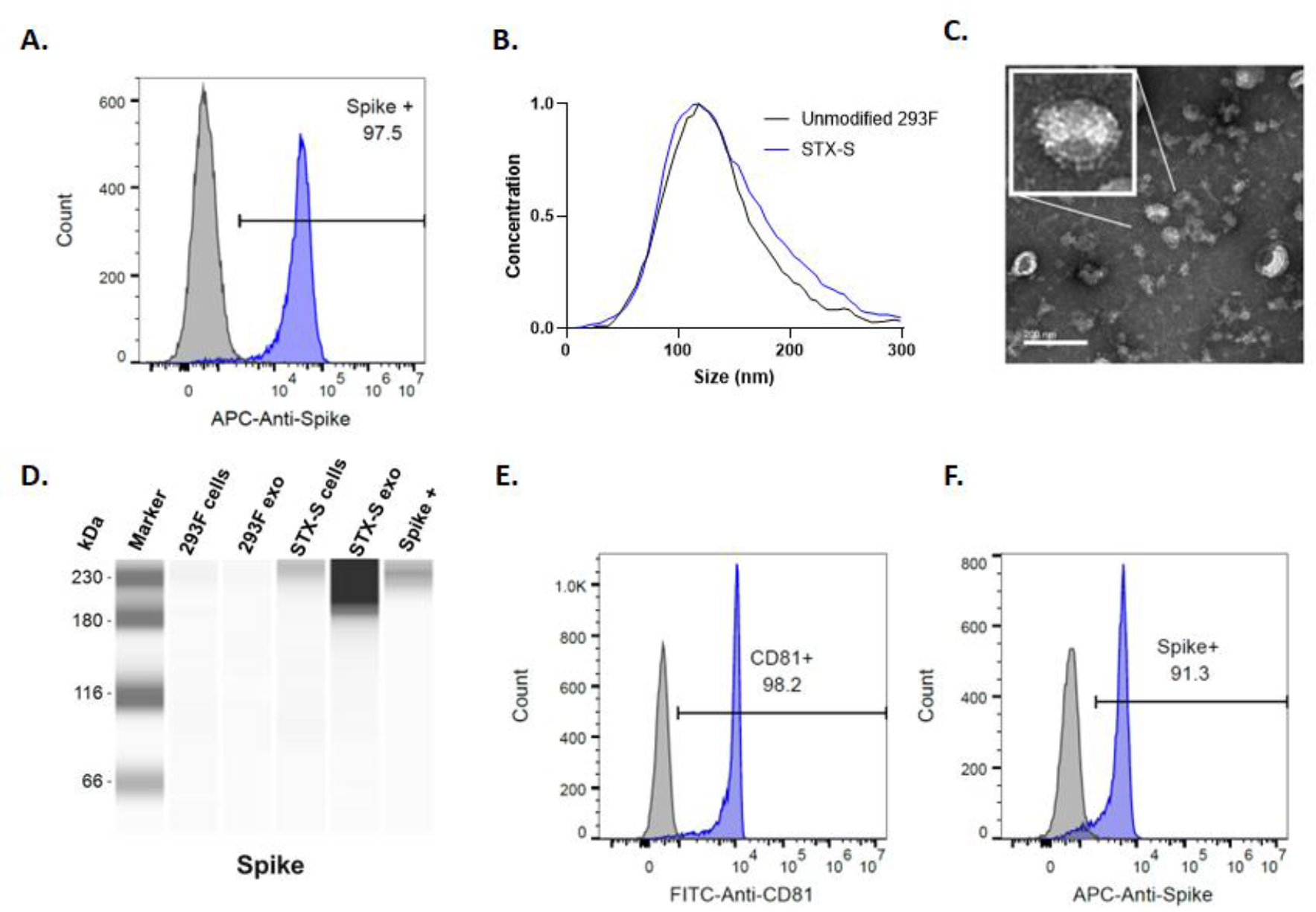
Characterization of STX-S cells and exosomes expressing spike. **A**. High expression of spike was detected on cell surface (blue) by flow cytometry. Parent, non-engineered 293F cells (grey) do not express SARS-CoV-2 spike protein, as expected (not shown). **B**. Size distribution of STX-S exosomes by ZetaView nanoparticle tracking analysis (NTA) compared to unmodified exosomes showed no change in distribution. **C**. TEM image of purified STX-S exosomes showed spike protrusions on the exosome membrane. Scale bar 200nm. **D**. Enrichment of Spike protein in exosomes was confirmed by Jess-automated Western Blot (Lane 5: STX-S exo). Lane 1: marker, Lane 2: non-engineered 293F cells, Lane 3: non-engineered 293F exosomes, Lane 4: STX-S cells, Lane 6: Spike protein. **E**. CD81 was detected on STX-S exosomes (blue) by flow-cytometry using a bead-assay compared with no signal from isotype control antibody (grey). **F**. Spike was detected on STX-S exosomes (blue) but not on 293F parental exosomes (grey).

STX-S exosomes were purified from the engineered STX-S cell culture supernatant using both lab-scale and large-scale purification techniques. Importantly, large-scale purification methods for clinical use have been developed and allow isolation of engineered exosomes with the same biochemical and biophysical characteristics as lab-scale exosomes (e.g., size distribution, particle concentration, surface marker expression, protein and lipid content) without contamination from cell components and ensure the separation of exosomes from proteins fractions in solution (data not shown). The ability to produce STX-S exosomes using a scalable process is critical for the clinical path of the STX-S vaccine.A Purified STX-S exosomes had an average concentration of 1.55E12 particles/mL with an expected, average diameter of 144.6 nm and an expected polydispersity index (PDI) of 0.152 (Fig. 2B). Note that a PDI below 0.2 is acceptable for nanoparticle delivery applications and is considered monodisperse [14]. STX-S exosomes were also analyzed by TEM imaging. As shown in Figure 2C, typical exosome size and morphology were observed with round, smooth nano particles detected that had a visible lipid bilayer. Importantly, spike protrusions were visible on the surface of the STX-S exosomes indicating the presence of spike protein on exosomes.

Spike protein expression was confirmed on STX-S cell lysates and exosomes using Protein Simple’s Jess automated Western blot, with enrichment of spike protein in the exosome samples (Fig. 2D). Spike expression was not detected in control, non-engineered 293F cells or control 293F exosomes as expected.

To demonstrate expression of spike on the exosome surface, a bead-based CD81 assay was used. This assay showed that >90% of STX-S exosomes bound to CD81 beads expressed spike on the exosome surface. In addition, the same bead-based assay but with a CD81 detection antibody was used to demonstrate that >95% of STX-S exosomes bound to beads expressed the characteristic exosome marker, CD81[15]. This assay demonstrates that the spike protein measured by Western blot is exosome-bound spike as it is expressed on the same particle as CD81 (Fig. 2 E, F).

The concentration of spike antigen in STX-S exosomes was then quantified by ELISA and 254 ng of Spike (average) was detected per 1E12 exosomes. Pre-clinical animal studies described below demonstrate that a range of 0.32ng to 33ng of the STX-S vaccine (per injection) was sufficient to induce a strong immune-response at both the humoral and cellular level in mice. Compared to other recombinant protein vaccines, the STX-S vaccine elicits a strong immunization using <1000X protein (ng of STX-S vs µg of traditional recombinant protein vaccines).

### STX-S vaccine induces robust production of SARS-CoV2-Spike specific IgG

The STX-S exosome vaccine was administered at varying doses into 8-10 weeks old female mice by either a single intramuscular (IM) injection or two IM injections. A second IM injection (referred to as boost injection) was delivered after a 3-week interval. Doses used across studies include: 0.32 ng, 3 ng, 10 ng, and 33 ng. PBS was used as a control in all studies.

The STX-S vaccine induced dose-dependent spike antibody production 2 weeks after the first prime injection in all animals in multiple studies. STX-S primed mice produced 8-fold (0.32ng dose) to greater than 30-fold (doses ≥10ng) greater SARS-CoV-2 Spike-specific IgG than control mice (Fig. 3 A, B, and data not shown). At day 35, two weeks post boost injection, robust spike antibody production was observed in all animals regardless of dose in all studies (>3 independent studies using >3 independent STX-S lots performed to date, with representative data shown in Figure 3). An increase of IgG against spike was observed ranging from 1000-fold at the lowest dose of 0.32 ng to over 2300-fold at the highest (≥10 ng). No significant difference was observed among the highest dose groups (10 ng and33 ng), suggesting that a dose higher than 3 ng of spike delivered by STX-S exosomes is sufficient to induce a strong spike antibody response in mice (Fig. 3 C, D).

**Figure 3.**
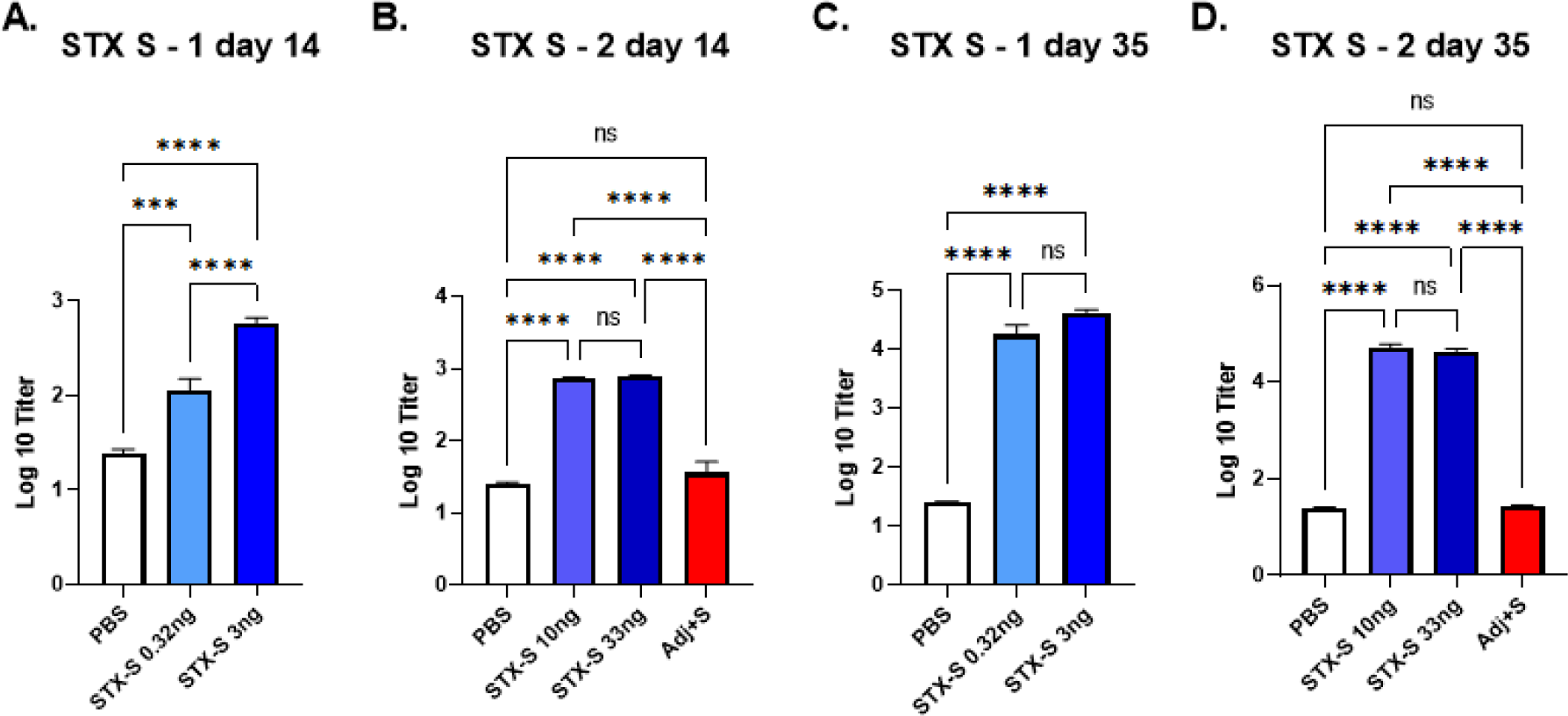
STX-S exosome vaccine elicited a robust immune response using significantly lower amounts of antigen compared to recombinant protein vaccines. The STX-S vaccine induced robust expression of SARS-CoV-2 spike antibodies in mice after 1 (day 14) and 2 (day 35) IM injections as analyzed by ELISA. PBS was used as a vehicle control in all studies. STX-S vaccine exosomes were administered at the doses described in methods and shown in representative graphs above. **A**. STX-S study 1, Day 14 after a single IM injection **B**. STX-S study 2, Day 14 after a single IM injection **C**. STX-S study 1, D ay 35 after complete immunization (two IM injections). **D**. STX-S study 2, Day 35 after complete immunization (two IM injections). Spike protein at the same dose in combination with adjuvant (Adj+S) was evaluated in STX-S study 2 (red bar) and did not elicit an immune response (not statistically significant from pbs control). N= 10/experimental group. Data are shown as mean ± SEM. **** p<0.0005, ***p<0.001, **p<0.01, *p<0.05, ns= not significant, 1-way ANOVA

Given the potent humoral response observed in all animals in multiple mouse studies using only nanograms of spike in STX-S exosomes, an additional group was added to compare the STX-S vaccine with a more traditional approach using a recombinant spike protein plus adjuvant-based vaccine. Recombinant spike protein was diluted to 33 ng and formulated with a commercially available adjuvant (Alhydrogel, Invivogen, Fig. 3 B, D, Adj +S in STX study 2 data, red bars) and injected into mice. The STX-S vaccine was injected in the same study using 10 ng and 33 ng spike delivered by exosomes for comparison (no adjuvant). PBS was used as a control. Blood collected 2 weeks after the boost injection (2^nd^ injection) showed that the STX-S vaccine elicited robust spike antibody production in all animals, as expected based on previous studies. Spike protein in combination with adjuvant at 33 ng was not statistically different than the PBS control and thus was not sufficient to elicit antibody production (Fig. 3 B, D). These data clearly demonstrate that the STX-S vaccine elicits a strong immunization by using significantly less protein, without the addition of an adjuvant, which is superior to the current mRNA-LNP and traditional recombinant protein vaccines.

### Potent neutralization induced by STX-S exosome injection

Immunization to the STX-S vaccine was further evaluated by assessing neutralizing antibodies against SARS-CoV-2 Delta variant (Fig. 4 A). Potent neutralizing activity was elicited using nanogram amounts of spike protein delivered by the STX-S vaccine in all analyzed animals in multiple studies. Mice from the 3 ng dose at day 40 and from the 10 ng dose at day 14 and 40 were tested for neutralizing antibodies against SARS-CoV-2 Delta variant (B.1.617.2, Fig 4A). The STX-S vaccine resulted in dose-dependent neutralization of the virus, estimated by the ability to protect infected cells from the virus-induced cytopathic effect (CPE).

**Figure 4.**
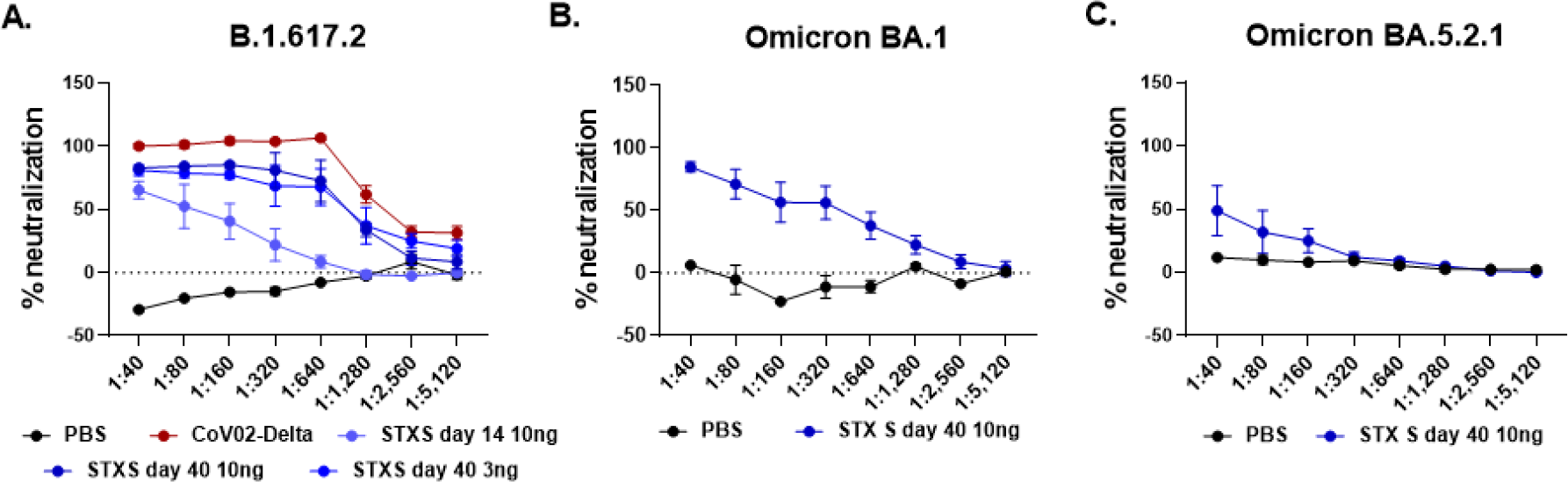
Potent Production of neutralizing antibodies after STX-S vaccination in mice using only 3 and 10 ng. **A**. STX-S vaccine generated potent neutralization against SARS-CoV-2 delta spike (B.1.617.2) **B**. STX-S vaccine resulted in strong neutralization of SARS-CoV-2 spike Omicron BA.1 **C**. STX-S vaccine resulted in neutralization of SARS-CoV-2 spike Omicron BA.5.2.1. Data are shown as mean ± SEM. COV-02-Delta = plasma from a patient immunized with Moderna’s mRNA vaccine with a breakthrough SARS-CoV-2 delta spike infection. N= 5/experimental group. Data shown as mean±SEM.

As shown in Fig. 4 A, the STX-S exosome vaccine induced neutralizing antibodies against delta spike by day 14 (after a single IM injection). By day 40, ∼3 weeks post STX-S boost, robust neutralization was observed in all animals regardless of dose. It is worth noting that one injection with 10 ng of spike by STX-S delivered a 64-78% neutralization against delta spike. Moreover, full immunization (2 IM injections, 3-10 ng of spike per injection) resulted in 91-98% neutralization against delta spike.

Importantly, the STX-S vaccine in mice induces a response comparable (i.e, not statistically different) to the human control plasma (COV-02, plasma from a patient fully immunized with Moderna’s mRNA vaccine, with breakthrough delta infection), with a complete neutralizing response with a dilution factor below 1:320 using 3 and 10 ng of spike by STX-S on day 40, Fig. 4 A).

Additionally, samples from the STX-S vaccine from the 10 ng dose on day 40 were tested for neutralizing antibodies against SARS-CoV-2 Omicron variants (Omicron BA.1 and BA.5.2.1). As shown in Fig. 4 B-C, a strong cross neutralization was observed for the STX-S treated mice (2 IM injections, 10 ng of delta spike per injection delivered by STX-S exosomes), achieving 74-97% (range, average of 84%) neutralization for Omicron BA.1 (Fig. 4 B, average of samples shown in graph) and 16-97% neutralization for Omicron BA.5 (Fig. 4 C, average of samples shown in the graph). Importantly, plasma from a patient immunized with a current leading mRNA vaccine (COV-02, collected 2 weeks after second injection) was used as an assay control and did not result in neutralization in these assays. These data show that protein-based vaccines, including the STX-S vaccine where nanogram amounts of spike protein are delivered by exosomes, result in broader protection against SARS-COV-2 variants.

### STX-S vaccine elicits spike-specific T cell responses

To characterize the T cell response to the STX-S vaccine, antigen-specific T cell responses were measured by ELISpot (Fig. 5). Vaccination with STX-S elicited multi-functional, antigen-specific T cell responses in all animals at both day 14 and day 40. Splenocytes were isolated from animals at day 14 (2 weeks after prime (1^st^) injection) and day 40 (∼3 weeks after boost (2^nd^) injection) and evaluated using ELISpot plates precoated with interleukin-4 (IL-4) or interferon-gamma (IFNγ. PBS was used as controls in the study. Baseline expression was compared to stimulation with 10 µg/ml of Spike protein (AcroBiosystem).

**Figure 5.**
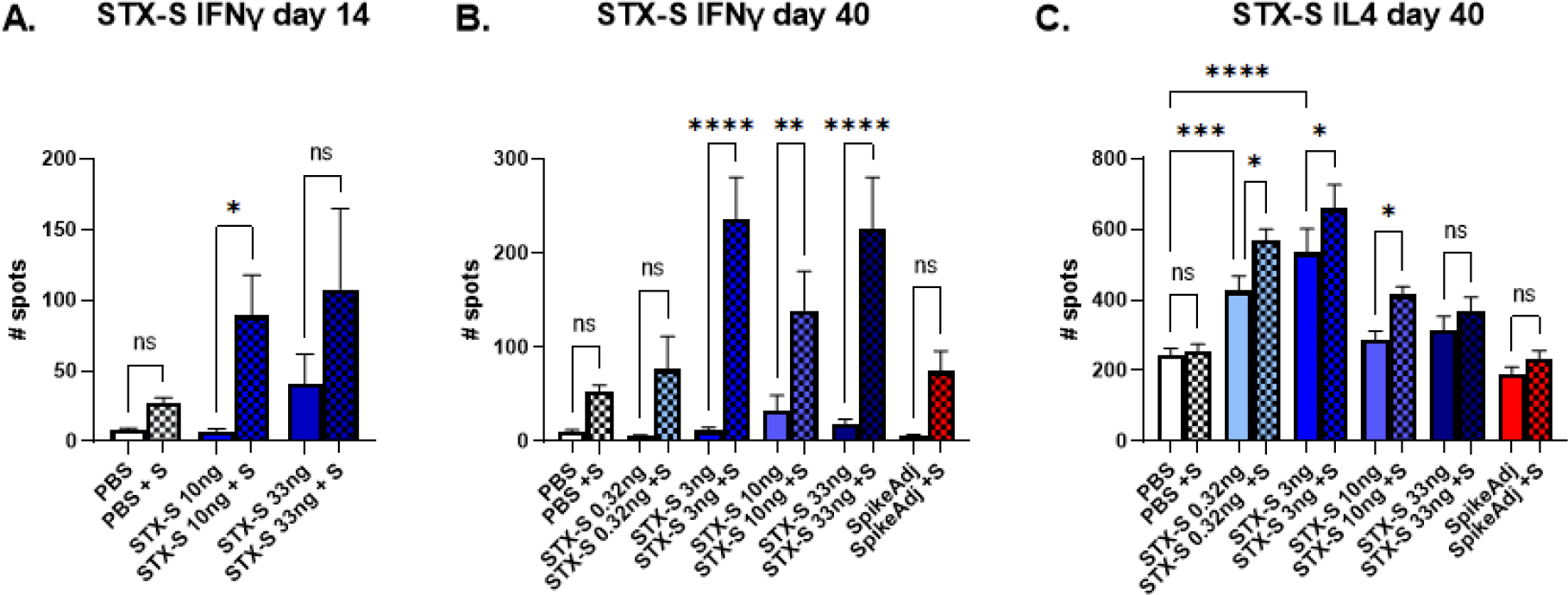
T cell mediated immune response: IFNγ and IL4. **A, B**. Strong IFNγ mediated T cell immune-response at day 14 (A) and day 40 (B). STX-S immunization resulted in increased IFNγ response as early as 2 weeks after the first dose after in vitro stimulation with 10ug/mL spike protein. **C**. IL4 mediated T cell immune-response at day 40. STX-S immunization resulted in increased baseline IL4 response compared to PBS control likely due to the time point of data collection, with an additional increase after in vitro spike stimulation. N= 10/experimental group. Data are shown as mean±SEM. *p<0.05, **p<0.01, ***p<0.005, ****p<0.001, 1-way ANOVA; ns= not significant.

Evaluation of IFNγ secreting cells in response to ex vivo spike stimulation showed a 10-fold increase in spleens immunized with STX-S by day 14 (p<0.05, Fig 5 A). Baseline IFNγ response was comparable between groups at day 14. A similar response was observed at day 40, with a statistically significant increase in IFNγ production by ∼8-fold at the lowest dose (p<0.05) and 12.6-fold (p<0.001) for mice receiving doses higher than 10ng (Fig. 5 B), suggesting a strong Th1-biased CD8+ T cell response.

Additionally, Type 2 IL-4 response was evaluated by ELISpot. At day 14, splenocytes from STX-S immunized mice showed an increase in IL-4 response at baseline (by ∼2-fold, p<0.05), with no additional increase after treatment with Spike protein (not shown). However, at day 40, splenocytes from STX-S immunized mice showed an increase in IL-4 response compared to PBS control group at baseline and after stimulation with spike protein, suggesting an increased Th2-biased CD4+ T cell response (Fig. 5 C) that will sustain the IgG production by B-cells. Together, these data demonstrate that the STX-S vaccine elicited a multi-functional, antigen-specific T cell response and induced a potent humoral immune response as demonstrated by high levels of neutralizing antibody not only against the delta SARS-CoV-2 virus but also the Omicron variant in all animals across multiple studies.

### STX-S exosome vaccine has no adverse health effects

Following vaccination with the STX-S vaccine, weight and general health were monitored throughout each study and no adverse effects of STX-S exosome administration were observed in any animal in any study (Supplemental Fig. 1). Moreover, complete blood count analysis was performed at the terminal timepoint (STX-S study 2) and no alteration of white or red blood cells was recorded (Supplemental Fig. 1). Pathological analysis of the STX-S vaccine at the highest dose of 33ng on day 14 (after a single injection) and on day 40 (after 2 injections) did not show any discernable alteration in ten tissues analyzed in any of the mice (Fig. 6). This safety profile was expected given that the STX-S vaccine does not utilize LNPs or an adjuvant and is dosed at ∼1/1000^th^ the dose of recombinant protein vaccines. Collectively, these data suggest that the STX-S vaccine elicits a potent immune response without adverse effects.

**Figure 6.**
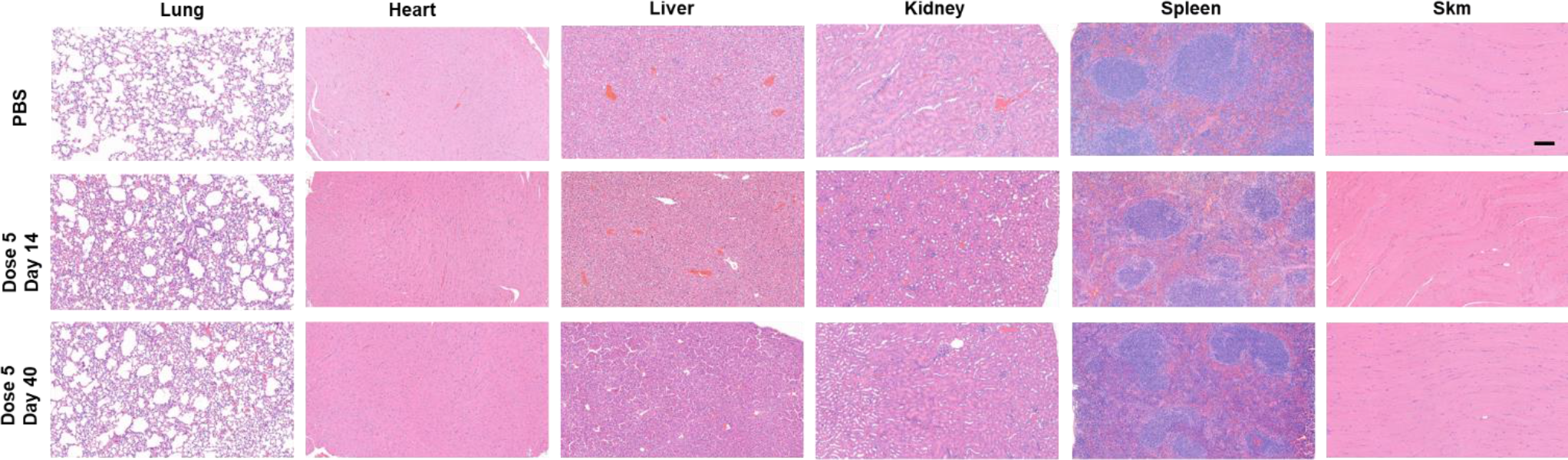
No adverse effects after STX-S administration. **A**. Representative images of H&E. All tissues were imaged at the same magnification. No adverse effect was observed after STX-S vaccine administration. Scale bar= 100um. SKM= skeletal muscle. N= 10/experimental group.

## DISCUSSION

Exosomes have gained great interest by the scientific community throughout the years for their high bioavailability, exceptional biocompatibility, and low immunogenicity, identifying them as promising drug delivery candidates [16]. Exosomes can cross biological barriers via their lipid membrane and endogenously expressed cargos and thus are able to effectively deliver engineered cargos as well [10]. As an endogenous product, highly produced by all cell types, exosomes are less prone to induce an adverse response thereby increasing safety. Together with their high stability at physiological pH and temperature, these unique exosome advantages make them an ideal delivery system for vaccinology.

Here, exosomes were engineered to express a target of interest on their surface. Engineered exosomes expressing native SARS-CoV-2 Spike (Delta variant), formulated as the STX-S vaccine, were delivered to mice by intramuscular injection according to a two-dose schedule with an injection interval at 21 days. A strong immune response at both the humoral and cellular level was induced as early as two weeks after the first injection, resulting in a potent antibody production and establishment of a cellular memory response after a complete immunization cycle (Fig. 3-5). IFNγ, mainly secreted by T helper 1 (Th1) cells, is responsible for cell-mediated immunity, while IL4, mainly secreted by T helper 2 (Th2) cells, induces significant antibody production [17]. Previous studies [18-21] showed that exosome mediated immunization induces a strong Th1 response, in accordance with our data. Note that Th1 responses may be higher in the BALB/c mouse strain and sex used in many vaccination studies, including the study presented here [22]. A Th2 immune response was also induced after STX-S vaccination, in accordance with the increased production of IgG and in particular, strong neutralizing antibodies. Given a cellular immune response is critical for virus control and clearance in acute infection, STX-S poses itself as a great candidate for efficient and long-lasting protection, able to induce both potent neutralizing antibodies and cellular responses.

The STX-S vaccine induced neutralizing antibodies against both Delta and Omicron spike variants: this clearly demonstrates the broad efficacy of the STX-S vaccine and suggests that both variants share epitopes important for the generation of neutralizing antibodies. Kumar et. al. reported that the Omicron variant had a greater affinity for the ACE2 receptor than the Delta variant because of additional mutations in the SARS-CoV-2 receptor-binding domain (RBD), indicating a higher potential for transmission [23]. The authors also reported common mutations in the N-terminal (G142D), the receptor binding domain (RBD, T478K) and the subdomain S2 (SD2, D614G, P681H) [23]. These mutations make both Delta and Omicron variants different from the original Wuhan wild-type SARS-CoV-2 virus. Indeed, the original mRNA vaccines designed against the Wuhan SARS-Cov-2 virus have offered little protection to the Delta and Omicron variants. Newer mRNA vaccines targeting these variants are being deployed but remain limited due lack of broad immunity and brief antibody response. Importantly, protein-based vaccines have been shown to have broader protection against mutating spike variants. Novavax recently reported that three doses of their NVX-COV2373 vaccine candidate was able to induce a potent neutralization against omicron subvariants[24]. This cross-variant neutralization has been further demonstrated here by our STX-S protein-based vaccine, after one single boost injection.

Few studies have reported using exosomes as a delivery system for vaccination [18, 19, 21, 25, 26]. In agreement with our results, a strong CD8+ T cell response was induced when those vaccine candidates were delivered by exosomes. Additionally, data from this study showed a potent B cell response, as demonstrated by IgG production and neutralizing antibodies. These results are likely due to the role of exosomes in natural intercellular communication and antigen presentation. In particular, numerous copies of the native Delta spike protein are likely present on the exosome surface, facilitating crosslinking to B-cell receptors. Moreover, viral proteins in exosome-based vaccines, including STX-S, could indirectly activate B cells and CD8+ T cells through this antigen cross-presentation. Kuate et. al. previously demonstrated that exosomes could be used for immunization of SARS-CoV-2. The authors replaced the cytoplasmic and transmembrane domains of spike with those of the G protein of vesicular stomatitis virus [26], allowing the incorporation of the protein into the exosome membrane. This exosome-based vaccine required two injections to induce a strong immune response, but both the number of exosomes and the concentration of spike needed for the immune response were unclear. While a direct comparison cannot be made, our STX-S vaccine was able to induce both higher antibody titer, and most importantly, efficient neutralization without modification in the spike sequence, therefore conserving the viral conformation.

The STX-S vaccine clearly presents multiple advantages over the currently available vaccines. First, the STX-S vaccine delivers the antigen through an entirely endogenous, autologous lipid bilayer, that can be easily integrated into the host cell membrane and facilitate engineered antigen presentation to immune cells. Membrane bound antigen can easily be presented to the circulating immune cells to quicky activate a response, while free antigen contained inside the exosome can additionally be processed by the lysosomal system and activate the cytotoxic T-lymphocyte response. Thus, utilization of a natural delivery system promotes efficient delivery and response compared with synthetic lipid nanoparticle technology. Second, the STX-S vaccine enables delivery of the native, trimeric, fully folded, and conformationally correct spike protein, which is likely to boost and broaden the immune response. Utilization of the native spike proteins allows the STX-S vaccine to skip the mRNA translation step, making the antigen readily available for cellular delivery and immune response. Incomplete translation and/or incorrect folding of the mRNA encoding the spike protein limit the amount of available antigen after vaccination, making the immune response extremely variable, with reduced efficacy as has been observed with mRNA vaccines. Third, the STX-S vaccine can be readily available and accessible for widespread global usage due to both known protein and exosome stability; exosomes are highly stable at physiological pH and temperature. Current mRNA COVID vaccines have a short shelf life and require preservation at extremely low storage temperatures. While additional studies will be required, the STX-S vaccine could be stored at 4°C for longer times or likely lyophilized for global vaccination.

Fourth, the STX-S vaccine does not require an adjuvant or synthetic lipid nanoparticle (LNP) for delivery and immune response. Protein-based vaccines require adjuvants (such as aluminum salt and squalene oil-in-water emulsion systems, MF59 (Novartis) and AS03 (GlaxoSmithKline), respectively [27]) to augment and orchestrate immune responses and influence affinity, specificity, magnitude and functional profile of B and T cell responses. The same Novavax vaccine uses a saponin-based Matrix-M adjuvant[28]. Current mRNA vaccines do not use an adjuvant, which combined with the need for mRNA translation prior to antigen availability, may also reduce long term efficacy. Approved COVID vaccines use LNPs to deliver the mRNA or adjuvants for the proteins with some side effects reported and long-term, repeat exposure effects unknown. Thus, there is an inherent safety benefit by eliminating these excipients (i.e., adjuvant, LNPs) in the STX-S vaccine. Additionally, pre-clinical safety studies previously performed in rabbits confirmed that Capricor’s exosomes are safe (data not shown). Together these data, along with exosomes natural delivery of the spike protein without the use of an adjuvant or synthetic LNP further enhance the safety profile of this vaccine.

Remarkably, the novel study presented here showed that exosome mediated antigen delivery can be achieved using nanogram amounts of protein without adjuvant, which has not been reported to date using any current vaccine. Inoculation using the same amount of spike antigen resulted in a statistically significant, strong immune response (2300-fold above PBS control, p<0.001) using the STX-S vaccine (no adjuvant) compared to no significant response using recombinant protein plus adjuvant groups (Fig. 3).

Approved recombinant protein-based vaccines use approximately 5-25 µg of antigen in conjugation with adjuvants to induce immunization [29-31]. Our novel data showed that the STX-S vaccine allows complete immunization correlated with high antibody levels, strong viral neutralization and both B and T cell memory with <1/1000^th^ of the spike protein administered in approved recombinant protein-based vaccines.

The broader neutralization of SARS-CoV-2 observed using only nanogram amounts of the native, delta spike antigen expressed by exosomes in this study opens the door to endless possibilities for vaccinology. Use of nanogram amounts of protein for vaccination enables extensive multiplexing of STX-based vaccines, which is not achievable by mRNA or recombinant protein-based vaccines using current technology. STX-based exosome vaccines can be rapidly engineered to express newly emerging variants, additional SARS-CoV-2 viral proteins, and other endemic viruses which can be delivered together in nanogram amounts in a multiplexed vaccine. Preliminary data in our hands suggest that STX-S vaccine can be combined with other viral proteins (i.e. nucleocapsid and H3N2 flu): STX-S immunogenicity is retained, and additional protection is gained by the vaccination with other viral antigens (data not shown). Rapid engineering, in a time frame similar to mRNA vaccines, coupled with our recent advances in exosome production, which rely on well-established cell engineering and particle purification technology, enables the STX-S vaccine technology to be rapidly deployed for numerous future vaccines. Indeed, preliminary data demonstrates the utility of a multiplexed STX vaccine.

While data from this study support the clinical development of the STX-S vaccine for immunization against COVID-19, it also opens the door to a whole new class of directed therapeutics outside of vaccines. STX exosomes could be engineered to selectively target organs, tissues of interest, or diseased cells to allow for safe and targeted delivery of compounds, RNAs, proteins, and peptides. For example, STX exosomes could be used to deliver chemotherapy by using a tissue-specific protein to target cancer cells with a chemotherapeutic loaded inside the exosome, potentially reducing the dosage and harmful side effects. Indeed, targeted therapeutics using exosomes are already in development.

In summary, the STX-S vaccine pre-clinical data presented here show that a potent B cell response was induced in immunized mice with less than 1 ng of spike protein displayed on exosomes, which was enough to elicit full neutralization of SARS-CoV-2 delta variant. Amazingly, neutralization of the Omicron variant was also observed using only 3 ng of delta spike protein, which has not been found using human serum immunized with two doses of the current mRNA vaccine, suggesting that our STX-S vaccine has a superior advantage as a broader vaccine candidate compared with current mRNA vaccines. The extremely low dose of spike protein (less than 1 ng dose compared with current recombinant protein vaccine dose at 5 ug or higher) without the use of an adjuvant or lipid nanoparticle, further enhances the safety profile of the STX-S vaccine for clinical development. Engineered exosome technology presented in this study has the ability to revolutionize the next generation of vaccines by providing a rapidly engineered, readily accessible vaccine which produces a potent, broader neutralization of the target virus, a robust T cell response, enhanced safety profile, and unlimited multi-valent vaccine cocktailing power by naturally delivering an extremely low dose of antigen via exosomes.

## Author contribution statements

MC contributed to the design, implementation, and analysis of all research in this paper and to the writing of the manuscript. MS did all the protein engineering and design of all the constructs used in this work for exosome engineering. YL, JBN produced the cell lines, carried the in vitro experiments, and contributed to the writing of the manuscript. MJL, MY, JA produced the exosomes used in the study. KE, MS contributed to the design and review of studies and the writing of the manuscript. MC, JBN, YL, CM, MJL, JA, MY, SYA, EC, RT carried out the experiments.

---

**Supplementary figure 1.**
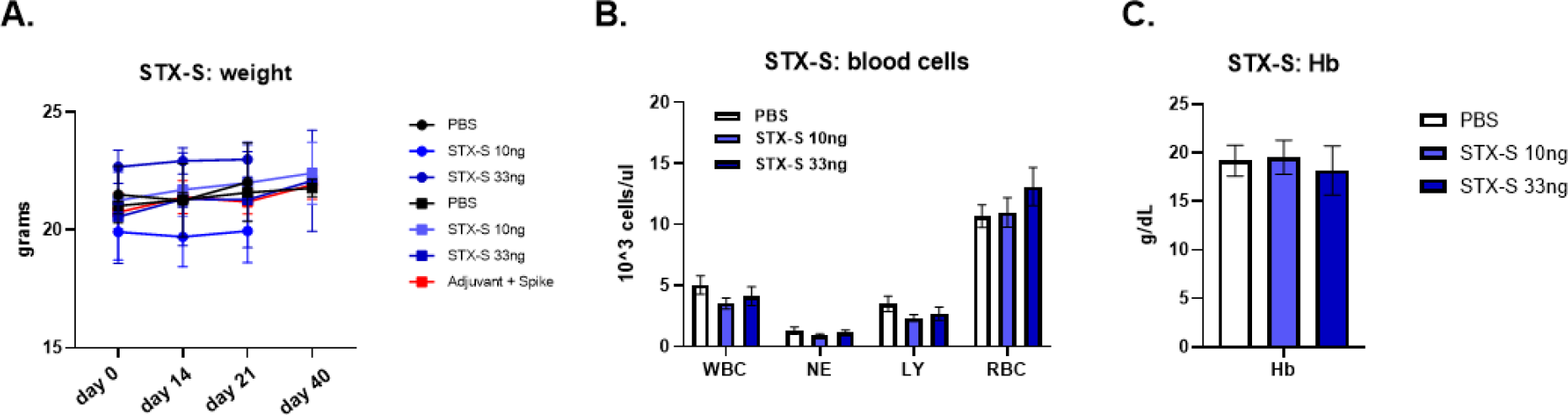
No adverse effects on complete blood count after STX-S administration. **A**. Weight progression at each timepoint. **B-C**. Complete blood count at terminal collection for mice from STX-S dose 4 and dose 5 WBC= white blood cells; NE=neutrophils; LY= lymphocytes; RBC= red blood cells; Hb= hemoglobin; SKM= skeletal muscle. N= 10/experimental group. Data are shown as mean±SE.

